# The protein repertoire in early vertebrate embryogenesis

**DOI:** 10.1101/571174

**Authors:** Leonid Peshkin, Alexander Lukyanov, Marian Kalocsay, Robert Michael Gage, DongZhuo Wang, Troy J. Pells, Kamran Karimi, Peter D. Vize, Martin Wühr, Marc W. Kirschner

## Abstract

We present an unprecedentedly comprehensive characterization of protein dynamics across early development in *Xenopus laevis*, available immediately via a convenient Web portal. This resource allows interrogation of the protein expression data in conjunction with other data modalities such as genome wide mRNA expression. This study provides detailed data for absolute levels of ∼14K unique *Xenopus* proteins representing homologues of ∼9K unique human genes – a rich resource for developmental biologists. The purpose of this manuscript is limited to presenting and releasing the data browser.

**Highlights:**

- Relative protein expression from stage IV oocyte, blastula, gastrula, neurula, and early organogenesis
- Biological triplicates with confidence intervals on protein expression reflect certainty in dynamic patterns
- Convenient time-series Web-browser integrated with the multi-media Xenbase portal
- Gene-symbol search and multi-gene protein/mRNA juxtaposition capabilities

## Genome-wide measurements of protein levels across key developmental stages

We profiled developmental stages (Nieuwkoop and Faber, 1994) spanning early development from stage VI oocyte and unfertilized egg (NF 0) through blastula (NF 9), gastrula (NF 12), neurula (NF 17 – 24) and tailbud (NF 30). Stage NF 24, for example, is characterized by the presence of blood islands and the first appearance of olfactory placodes. The last time point (NF 42) is taken long after the heartbeat has started and the tadpole has hatched, most of the cardio-vascular and digestive (liver, pancreas) system having formed. Our processing pipeline for quantitatively measuring levels of protein is as previously described (Gupta et al., 2018; Peshkin et al., 2015). Proteins were digested into peptides and change of abundance was measured by isobaric labeling followed by MultiNotch MS3 analysis (McAlister et al., 2014); absolute protein abundance was estimated via MS1 ion-current (Schwanhäusser et al., 2011; Wühr et al., 2014). Protein abundance levels were measured at ten key stages (oocyte VI, egg=NF 0, 9, 12, 17, 22, 24, 26, 30, 42). Our primary dataset is comprised of 14940 protein profiles. We collected and profiled the data in three independent biological replicates, but for the purposes of presenting the data at Xenbase, combined all information using our “BACIQ” pipeline (Peshkin et al., 2019) which produces the most likely patterns of relative protein abundance and confidence intervals for these.

## Xenbase visualization

Our data is integrated into Xenbase via a convenient Web portal.

Users can get to a given gene’s protein expression graph by clicking the “Proteomics” link on Xenbase’s corresponding gene page (e.g. *kank2*’s gene page is at www.xenbase.org/gene/showgene.do?method=display&geneId=997066). Since *Xenopus laevis* is a pseudotetraploid where many genes exist as two homeologues “L” and “S”, there are separate links for L and S homeologs at each gene page. If expression data do not exist for a gene, the user is notified with a “No data” message at the graph page. In order to compare protein to mRNA expression at the same graph, the user can turn on the mRNA profile by clicking on the mRNA checkbox. The graph’s X axis represents the oocyte and embryonic developmental stages, scaled per hours lapsed between the stages after fertilization. Protein data are displayed for Nieuwkoop-Faber stages: oocyte VI, egg, NF-9, NF-12, NF-17, NF-20, NF-22, NF-24, NF-29-30, and NF-42. mRNA data, if present in the database, are displayed for stages oocyte VI, egg, NF-8, NF-9, NF-10, NF-12, NF-15, NF-20, NF-25, NF-29-30, NF-35-36, and NF-40. For proteins, the left Y axis shows the relative protein expression levels. This represents the decimal fraction, at a given stage, of total protein agglomerated over all profiled stages. In other words, if all stages contain the same protein level, a user would see a straight line at 0.1 (10 samples of 10 respective stages at 10% each). When the mRNA display is on, the graph shows mRNA expression levels in Transcripts per Million (TPM) units on an auxiliary Y axis on the right. As new genes are added or removed, units on both Y axes are rescaled to accommodate the display of all selected protein and mRNA profiles within the display area. Hovering the mouse pointer over a data point, represented as a filled circle, shows the corresponding data value. These expression values are shown by the plotting routine verbatim, without any pre- or post-processing.

Protein identifiers generated from our experiments were tagged with JGI laevis genome 9.1 identifiers. These were mapped to gene models in Xenbase to determine the corresponding gene symbols. Out of the total 14,941 original proteins 12,218 were matched with a gene model and respective gene symbol. Using a symbol, such as *hk1*, to add profiles to the graph results in both L and S homeologs displayed, provided the data is available.

All profiles, including for genes not matched to a Xenbase gene symbol, can be added to an existing graph by entering the corresponding *X. laevis* 9.1 mRNA model identifiers into the search box. For example, Xelaev18034790m will show the profile for *hk1.L*. Note that gene symbols and model identifiers are case insensitive. It is possible to add multiple genes at the same time by entering the corresponding gene symbols or model identifiers as a comma separated list.

By default, the graph includes protein expression levels and the corresponding error bars reflecting 90% confidence intervals. Random colors are chosen each time the graph page is loaded. Reloading of the page via the browser window will trigger the use of different colors. Hovering the mouse over highlighted data points for either protein or mRNA graphs will display the corresponding expression values. Users can click on the Redraw button to reset the graph, which involves removing the expression values, selecting the display of error bars, and unselecting the display of the mRNA graph. Saving the graph as an SVG file is also supported, and the captured image will contain all the elements displayed at the save time.

Following each gene symbol or model identifier the number of peptides used to characterize each protein expression is given in parentheses. Naturally the more peptides, agreeing among themselves there are the tighter confidence intervals would come out.

Code and database operations behind the visualization are fast and efficient. We use D3.js, a versatile Javascript-based graphing tool, to draw the graphs. Protein data are stored as comma-separated values (CSV) in the database. Each entry in the database consists of a gene symbol and a series of expression values corresponding to the represented developmental stages. The CSV data are encapsulated in a JSON object before being passed on to the visualization code. mRNA data are also transferred as JSON objects.

## Overview of protein dynamics patterns

Most developmental studies have focused on genes which are variable, i.e. expressed at different times, places and circumstances. What is not clear is whether these are exceptional cases or whether embryos are constantly changing the majority of proteins. It has been noticed before that in *X. laevis* there is little new protein synthesis from fertilization up to neurulation (Lee et al., 1984). Overall protein synthesis does not change much throughout these periods and remains at approximately 100 ± 20 (sd.) ng per hour or about 0.4% per hour of the total non-yolk protein content. Proteins that appear stable throughout our experiment are therefore likely to be made early and not degraded, rather than maintain a constant level through high production rates and high turnover. MS analysis allows us to see which proteins are stable and which are dynamic, thought bulk measurements bias the interpretation toward the most abundant proteins.

Figure 2 presents six main temporal trends of relative protein abundance via the medians of clusters (K-means clustering using cosine distance as similarity measure). Please note that for this analysis all stages were normalized so that ribosomal proteins are flat. It is possible that ribosomal protein levels change significantly after stage ∼30. This would lead to a significant increase in total protein mass, which is not shown here. The thickness of the median line reflects the number of proteins that fall into the respective cluster. The largest cluster of 4550 proteins or 30% of all proteins is flat. The four next largest clusters together 10044 or 67% contain proteins whose abundances are growing with time more or less aggressively. Only 346 or 2% of proteins are in the cluster of degradation. Clearly the more dynamic the trend, the fewer proteins fall into that category. This strongly confirms our previous observations (Peshkin et al., 2015). There are definitely dynamic patterns of proteins getting rapidly synthesized and degraded during development, but these are not sufficiently well represented to define a cluster of their own when all ∼15K proteins are forced to fall into six representative clusters.

**Figure 1:**
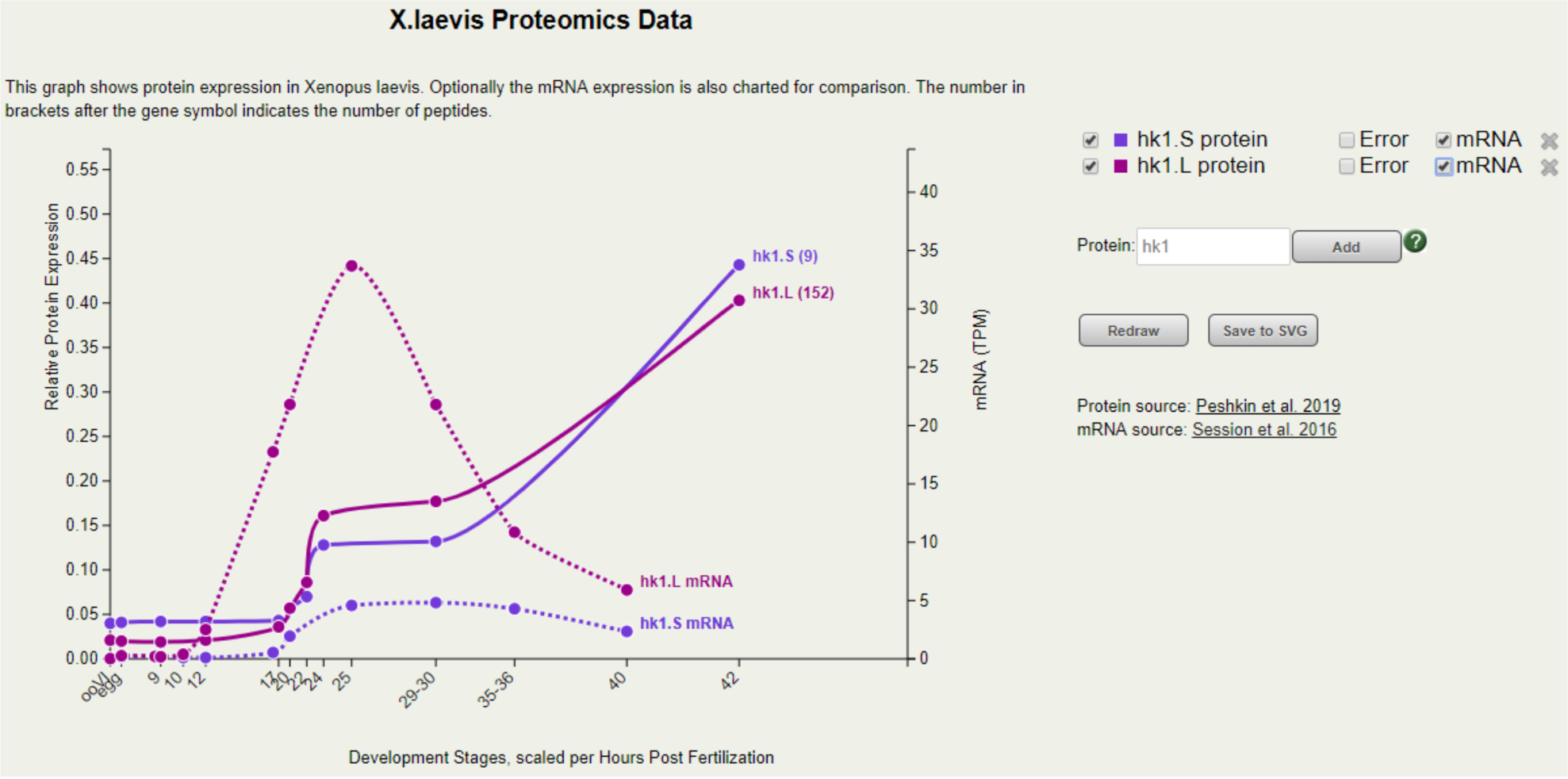
Screen capture of a browser window fragment showing jointly mRNA and protein expression for both L and S alleles of Hexokinase HK1 within Xenbase gene-centric view.

**Figure 2:**
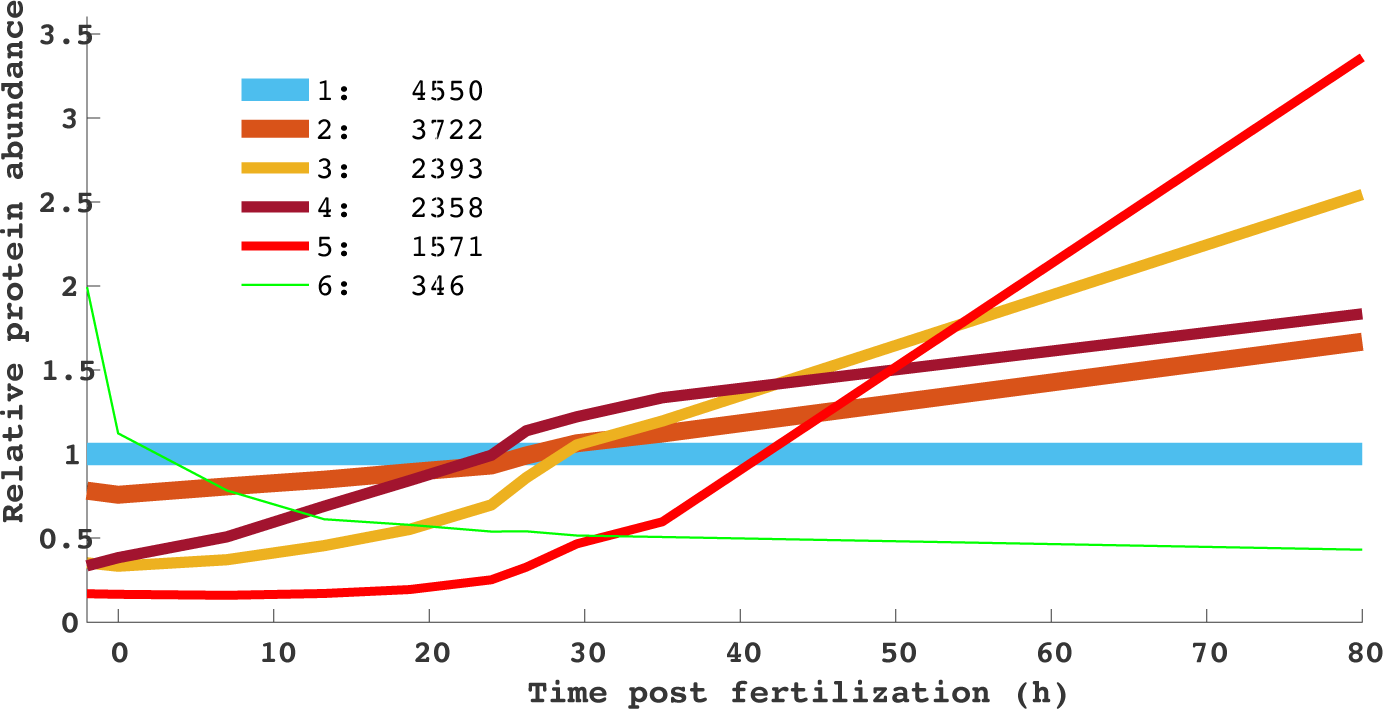
Key trends in dynamics of protein expression illustrated by mean patterns in six clusters resulting from K-means clustering based on cosine similarity distance.

### Protein measurements are reliable

We assessed the level of agreement across three biological replicates as described below and found it to be consistent, suggesting not only high quality of our measurements but also a high degree of control of the protein expression in embryogenesis.

We identified 32114 protein measurements that had at least one repeat across biological triplicates in our data and we asked how consistent the measurements are across replicates. To quantitate this concordance we calculated the similarity of the dynamic patterns by Pearson correlation coefficient (see histogram in Figure 3.left). In principle, an agreement between two measurements could be trivially explained by a common general trend, such as overall protein synthesis. To control for this we chose as a baseline the correlation between proteins matched uniformly at random across repeats (see gray histogram in Figure 3.left). Indeed there is some residual correlation in such randomly paired measurements, which is most easily explained by a general trend of proteins to be stable or increase in level with time, as shown in Figures 1 and 2. However, properly matched repeats (the green bars in Figure 3.left) show much more striking agreement. This agreement allows us to confidently mix together measurements from all three repeats in one united dataset which we release and present at Xenbase. This dataset presents more total proteins and more peptides per protein than each individual repeat. Detection of different proteins in repeated experiments from could be real, and be due to actual variability in the repertoire of proteins expressed in different animals. Alternatively, it could be explained by abundance bias, that is by that the more abundant a protein is, the more likely the mass spec instrument would detect its peptides.

**Figure 3:**
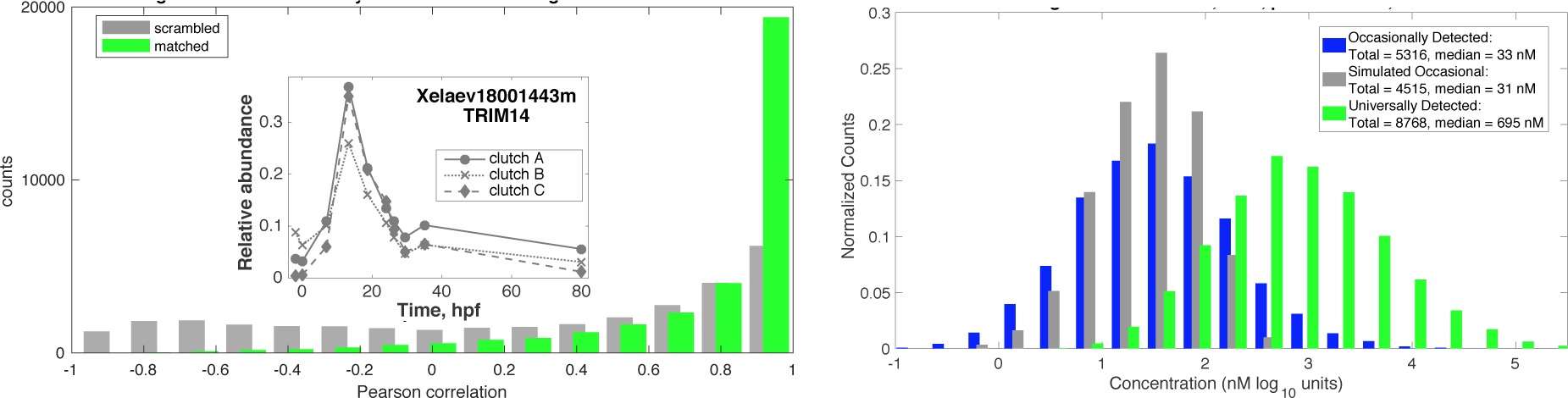
(left) Similarity between repeated measurements of the same protein is illustrated as a histogram of Pearson correlation between all 32114 repeated temporal patterns. As a baseline we present the same histogram for scrambled matches. The insert presents three repeated measurements of TRIM14 protein to illustrate the median level of agreement. (right) Concentration distributions for proteins detected in all three experimental repeats (green) compared to proteins missed in at least one repeat (blue) and simulated via concentration bias (gray).

To test the importance of abundance bias we compare the concentration distribution between proteins detected in all three independent proteomics mass spec experiments (“universally detected proteins”) with those that were not detected in at least one of the experiments (“occasionally detected proteins”) (see Figure 3.right). As expected the occasionally detected proteins (blue chart) are substantially less abundant compared to universally detected proteins (green chart): a median value of 30 nM as compared to 700 nM. In order to verify that this difference is explained by the abundance bias, we created a batch of “simulated occasional” proteins as follows: from a batch of all ∼15 000 observed proteins we sampled three simulated sub-sets each of a size of corresponding real clutches, namely: 9345, 11001 and 13451 proteins, and represented the concentration histogram of the proteins observed in at most twice out of three sub-sets (gray chart). Even though the Kolmogorov-Smirnov test rejects the null hypothesis (p = 2.9e-40) that the simulated occasional and the actual occasional concentration distributions are the same, they appear pretty close. The difference in turn can be explained by:

- a more complicated sampling bias in the mass spec instrument;
- actual biological differences in protein expression across the clutches;
- the actual underlying distribution of protein expression in the sample being much heavier in the low-abundance proteins than what we observe across our three replicated experiments, which means we are sampling in the simulation from an unrealistic overall abundance distribution.

## Confidence Intervals

All peptides across multiple repeats corresponding to a given protein are assembled and processed to reflect dynamics of that protein using our recently developed data analysis pipeline called BACIQ. The details of that method are outside of the scope of this manuscript, but we illustrate overall concept in Figure 4.left using three developmentally important genes. Specifically, Figure 4.left shows a sample panel from Xenbase.org displaying protein expression for three developmentally important genes. We illustrate variable degree of confidence in expression pattern as reflected by the shaded area of 90% confidence intervals. Number in parenthesis after each gene symbol indicates the number of peptides detected for respective protein. BACIQ confidence interval integrate the number of peptide measurements, the agreement across peptide measurements and the signal level of every peptide measurement. The general tendency for the confidence intervals is to get tighter with more peptides measured as long as peptides agree among themselves. Also peptides measured with higher signal to noise ratio bring more confidence than low signal peptides. It is possible, as illustrated in Figure 4.left to have higher confidence in one part of the interval and lower in another because peptides measurements have more agreement in one part of the trajectory.

**Figure 4:**
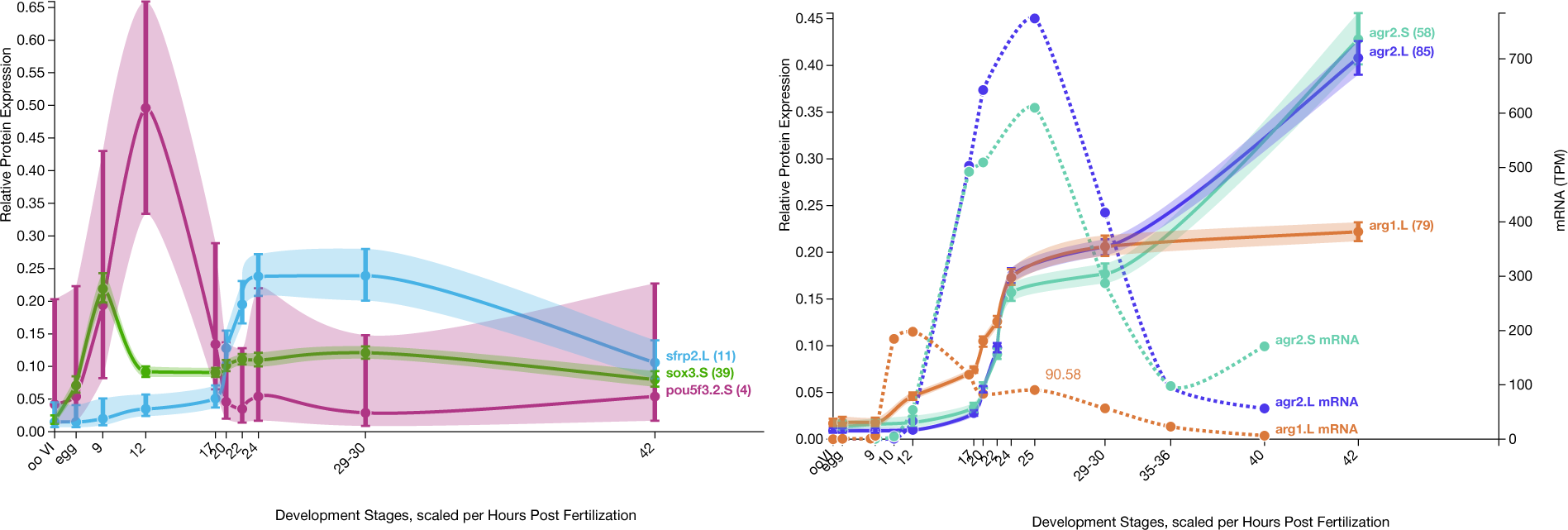
(left) Protein expression of three developmentally important genes with respective confidence intervals. (right) Examples of protein dynamics not readily explained by respective changes in mRNA expression.

### Discordance between mRNA and protein patterns

A master equation for a protein’s abundance would represent its accumulation to be proportional to the respective mRNA concentration minus the protein loss which is independent of the protein or RNA concentrations. Many proteins that we measured in the embryo follow this functional relation, though the first order rate constants for synthesis and the zero order rate constants of course differ for each protein (Peshkin et al 2015). Figure 4.right represents three of discordant cases where the levels of protein do not follow this simple pattern. This figure shows the accumulation of two homeologues of disulfide isomerases anterior gradient 2 (agr2) and in addition arginase 1 (agr1). Agr2 is in particular important for mucin secretion in the *Xenopus* cement gland where it was first discovered. Agr2 RNA and protein are rapidly synthesized after stage 17 at the time of the appearance of the cement gland and stop accumulating as the cement gland is fully formed. Sometime after stage 17 the RNA drops steadily as the protein increases slightly, a result counter to our expectations protein dynamics can not be explained by RNA. Subsequently the RNA levels drop to 10% of their original level while protein levels continue to climb, a second discrepancy. The protein data has high confidence, is virtually the same for the two isoforms and the pattern is also very similar for Arg1. Arginase catalyzes the hydrolysis of arginine to ornithine and urea. This kind of discrepancy was anticipated by the many early studies postulating translational control, where stored mRNA is supposed to be regulated post transcriptionally. What seem most interesting about such examples in this comprehensive data set is how relatively rare those are found.

## Discussion and Cautionary Remarks

In the interest of full disclosure we decided to spell out several possible issues which could result from hastily interpreting the protein dynamics plots.

The levels of all proteins in this round of data sharing were <u>normalized to the mean expression</u> <u>of ribosomal proteins</u>. If this normalization is not taken into account when interpreting the data. At the time of writing this manuscript we did not consider changes in the level of ribosome protein expression, particularly at the latest surveyed stages.

<u>Spline based curves</u> connecting data points should not be confused with the actual measurements. The time points making up developmental series are spaced unevenly, particularly the later time points are separated by tens of developmental hours. In displaying continuous curves we have made no attempt to perform biologically plausible interpolation.

For the purpose of reporting proteins we used both so-called “unique” and “razor” (non-unique) peptides. This allowed us to provide information for more proteins at the cost of aliasing the measurements across proteins with shared peptides. As a result this is not the best way to compare <u>expression of homeologues</u> since these share most if not all, peptides.

Note that the <u>number of peptides provided</u> with each protein measurement corresponds to distinct spectra-peptides measurements, which often include the same (in sequence) peptide being measured several times by mass spectra, as opposed to a number of peptides, unique in sequence. This could lead to confusion as many hundreds of peptides are indicated for what could be a rather short but abundant protein.

<u>Gene symbols</u> as displayed in Xenbase browser correspond to the names matched to Xenbase models. The gene models evolve with new releases of genome assembly and new rounds of assembly annotation. Gene symbol nomenclature is updated for other reasons. As a result one should use caution in interpreting the gene symbols verbatim. Our suggestion is to go back to the sequence of the original gene/protein used for mapping and re-do your own BLAST-based sequence homology analysis if you need to rely on gene symbol.

There are many <u>post-translational modification</u> (PTMs) which are not accounted for in our sample preparation, instrument and bioinformatics pipeline. Naturally, a modified peptide is detected differently from an un-modified one by the instrument. While it is in principle possible to hypothesize a PTM such as methionine oxidation or serine phosphorylation and search the spectra accordingly, controlling false-discovery rate for such searches is poorly understood at the moment. As a result any dynamic patterns which appear as relative protein abundance changes have to be taken with caution. Such patterns might result from changes in the level of some PTMs and in cases of where only a few peptides constitute the protein dynamic calculations, PTMs might strongly affect or entirely define the dynamic pattern.

<u>Relative protein abundance</u> of different proteins cannot be inferred from the plots. Plotting several proteins together might produce a graph where one of the proteins might appear more or less abundant than the other at a given time point. The only information meant to be displayed is that of dynamic pattern of expression of each protein relative to its own expression at a different time point.

Relatively <u>low levels of protein</u> are indistinguishable from zeros. The specifics of the used measurement methods in quantitative mass spectrometry applied in obtaining this dataset do not allow us to confidently measure the absence of protein in one sample which is present at substantial level in another one. E.g. see Pappireddi et al. for a review on methods and challenges for multiplexed proteomics (Pappireddi et al., 2019). Therefore it would not be correct to take our data as evidence for presence of low level rather than complete absence of a protein.

In Conclusion, we have focused this manuscript on releasing the protein dynamics plots in the early development of frog eggs from fertilization to long past hatching. Our exposition is aimed at presenting the data and providing several examples of data interrogation, accessible immediately via convenient Xenbase web portal. We assume Xenbase users will cite this manuscript as well as full paper containing in-depth analysis of protein expression patterns which is still in preparation.

## Author Contributions

Conceptualization: LP, MW, MWK; Methodology: LP, MW, MWK; Experiments: LP, MW, RMG; Software: LP, AL, DZW, TJP, KK; Formal Analysis: LP, AL, AMK; Resources: LP, MK, PV; Data Curation: LP, DZW, TJP, JG; Writing: LP, MWK; Supervision: LP, PV, MWK.

## Acknowledgements

LP and MK were supported by NIH grants R01HD091846 and R01HD073104. MW was supported by NIH grant R35-GM128813.

## Experimental Procedures

*Xenopus laevis* J-line embryos were collected according to NF system (Nieuwkoop and Faber, 1994) at stages oocyte VI, egg (NF0), 9, 12, 17, 22, 24, 26, 30 and 42. Embryos were de-jellied in 2% cysteine, pH 7.8, and flash frozen for later preparation. Stage VI oocytes were obtained by surgery from the same females as oocytes for fertilization.

MS sample preparation and data-analysis was performed essentially as previously described (Gupta et al., 2018; Sonnett et al., 2018; Wühr et al., 2015). Embryos were lysed and yolk removed via centrifugation (Wühr et al., 2014). Proteins were purified via methanol chloroform extraction (Wessel and Flügge, 1984), digested with LysC and labeled with ten-plex TMT. LC-MS experiments were performed on an Orbitrap Elite (Thermo Fischer Scientific) using the MultiNotch MS3 method (McAlister et al., 2014). For quantification we used both unique and non-unique peptides that matched the protein in the reference database. For the quantification of each protein we used a weighted sum of TMT Signal/FT-Noise intensities of its assigned peptides.

### Lysis

Frozen embryos/eggs at each stage were thawed and 5-6 uL of lysis buffer (250 mM sucrose (EMD cat# 8550), 10 mM EDTA (VWR cat# VW1474-01), 1 tablet Roche Complete mini protease inhibitor (cat# 11836153001) and 1 tablet Roche Phos STOP tablet (cat# 04906837001) per 10 mL, 25 mM HEPES (Sigma cat# H3375-500G) pH 7.2, 10 uM Combrestatin 4A (Santa Cruz cat# sc-204697), 10 uM Cytochalasin D (Santa Cruz cat# sc-201442), 1 mM TCEP (ThermoFisher cat# 20490) was added per embryo/egg on ice. The embryos/eggs were lysed by pipetting followed by vortexing well.

### Yolk removal

The yolk was removed from the samples by spinning at 4,000 xg for 4 minutes in a microcentrifuge. The lipids in the sample were then resuspended by lightly flicking the tube being careful not to resuspend any of the yolk. The supernatant was transferred to another tube and 2% SDS (Amresco cat# M112-500ML) was added. HEPES (Sigma cat# H3375-500G) (pH 7.2) was added to a final concentration of 100 mM.

### Alkylation and cysteine protection

DTT (5 mM, pH ∼8.0, Sigma cat# 43819-1G) was added to each sample and the sample was incubated at 60 C for 20 minutes, then cooled to RT. Next, NEM (15 mM, Sigma cat# E3876-25G) was added to each sample and the samples were incubated for 20 minutes at room temperature. Finally, DTT (5 mM, pH∼8.0, Sigma 43819-1G) was added again and the samples were incubated at room temperature for 10 minutes.

### Enzymatic Digestions

The proteins in the sample were precipitated using methanol/chloroform (Friedman, D.B. and Lilley K.S. “Quantitative proteomics for two-dimensional gels using difference gel electrophoresis (DIGE) technology” in John M Walker, Protein Protocols, 3rd Edition, Humana Press, 2009.) and were resuspended in 6 M Guanidine HCl (Sigma cat# G3272-1KG, buffered using 50 mM EPPS (Alfa Aesar cat# A13714), pH 9.0) to an estimated protein concentration of 5 mg/mL. The samples were heated to 60 C for 5 minutes and then allowed to cool to room temperature. The protein concentrations of each sample were determined by BCA assay (ThermoFisher cat# 23225). Next each sample was diluted to 2 M Guanidine HCl with 5 mM EPPS (pH 9.0, Alfa Aesar cat# A13714), ensuring that the pH was at least 8.5. Lys-C(Wako Chemicals cat# 129-02541) was added at the higher of 1:100 w/w or 20 ng/uL and incubated for 12 hours at room temperature. The samples were diluted to 0.5 M Guanidine HCl with 5 mM EPPS (pH 9.0, Alfa Aesar cat# A13714), ensuring that the pH was at least 8.5. Lys-C (Wako Chemicals cat# 129-02541) was added again as above and allowed to digest for at least 15 minutes at room temperature. Additionally, Trypsin (Promega cat# V5113) was added at the higher of 1:50 w/w or 10 ng/uL and allowed to digest at 37 C for 8 hours. Samples were speed-vacuumed to dryness.

### TMT labeling

Each sample (stage) was labeled using a distinct channel of TMT label (ThermoFisher cat# 90111 & A34807). Samples were resuspended in 500 mM EPPS (pH 8.0, Alfa Aesar cat# A13714), checking that the pH is close to 8.0, and incubated at 65 C for 5 minutes. 15 uL of TMT reagent (20 mg/mL, ThermoFisher cat# 90111 & A34807) were added per each 100 ug of protein and the reactions were incubated for 2 hours at room temperature. A small subset (∼1 ug/condition) was quenched and tested for TMT labeling efficiency, missed cleavage rate, and normalized total peptide count. Reactions were quenched by first heating the samples to 60 C for 5 minutes and then cooled to room temperature followed by incubation with 0.5% hydroxylamine (Sigma cat# H9876) for 15 minutes at room temperature. The samples were then combined in another tube containing phosphoric acid (JT Baker cat# B34P0200) that was at 5% of the total combined volume. The combined sample was then speed-vacuumed to dryness.

### Mass spectrometry analysis

Data were collected by a MultiNotchMS3 TMT method (McAlister et al., 2014) using Orbitrap Fusion mass spectrometers (Thermo Fisher Scientific) coupled to a Proxeon EASY-nLC 1000 liquid chromatography (LC) system (Thermo Fisher Scientific). The capillary column was packed with C18 resin (2.6 μm, 150 Å, Thermo Fisher Scientific). Peptides of each fraction were separated over 4 hour acidic acetonitrile gradients by LC prior to mass spectrometry (MS) injection. The first scan of the sequence was an MS1 spectrum (Orbitrap analysis; resolution 120,000; mass range 400−1400 Th). MS2 analysis followed collision-induced dissociation (CID, CE=35) with a maximum ion injection time of 150 ms and an isolation window of 0.7 Da. In order to obtain quantitative information, MS3 precursors were fragmented by high-energy collision-induced dissociation (HCD) and analyzed in the Orbitrap at a resolution of 50,000 at 200 Th. Further details on LC and MS parameters and settings used were described recently (Paulo et al., 2016a).

### Mapping and normalization

For mapping of protein data (peptide-Spectra matches for MS) we used as a main database of reference sequences *X. laevis* genome assembly (DoE JGI REF; v9r1 of assembly v1.8.3.2: a total of 45,099 sequences) downloaded from Xenbase (Bowes et al., 2010) ftp://ftp.xenbase.org/pub/Genomics/JGI/Xenla9.1/1.8.3.2/ Peptides were searched with a SEQUEST-based in-house software with a target decoy database strategy and a false discovery rate (FDR) of 2% set for peptide-spectrum matches following filtering by linear discriminant analysis (LDA) and a final collapsed protein-level FDR of 2%. Quantitative information on peptides was derived from MS3 scans. Quant tables were generated requiring an MS2 isolation specificity of >75% for each peptide and a sum of TMT s/n of >100 over all channels for any given peptide and exported as TAB-separated files. Details of the TMT intensity quantification method and further search parameters applied were described recently (Paulo et al., 2016b). The channels were further normalized assuming the ribosome expression is constant across developmental stages. Specifically a normalization factor was obtained by summing up signal for peptides belonging to any of the ribosomal proteins and dividing raw data by that factor. Peptides were summarized per protein using BACIQ pipeline (Peshkin et al., 2019).

## References

Bowes, J.B., Snyder, K.A., Segerdell, E., Jarabek, C.J., Azam, K., Zorn, A.M., and Vize, P.D. (2010). Xenbase: gene expression and improved integration. Nucleic Acids Res. 38, D607-612.

Gupta, M., Sonnett, M., Ryazanova, L., Presler, M., and Wühr, M. (2018). Quantitative Proteomics of Xenopus Embryos I, Sample Preparation. Methods Mol. Biol. Clifton NJ 1865, 175–194.

Lee, G., Hynes, R., and Kirschner, M. (1984). Temporal and spatial regulation of fibronectin in early Xenopus development. Cell 36, 729–740.

McAlister, G.C., Nusinow, D.P., Jedrychowski, M.P., Wühr, M., Huttlin, E.L., Erickson, B.K., Rad, R., Haas, W., and Gygi, S.P. (2014). MultiNotch MS3 enables accurate, sensitive, and multiplexed detection of differential expression across cancer cell line proteomes. Anal. Chem. 86, 7150–7158.

Nieuwkoop, P.D., and Faber, J. (1994). Normal table of Xenopus laevis (Daudin): a systematical and chronological survey of the development from the fertilized egg till the end of metamorphosis (New York: Garland Pub).

Pappireddi, N., Martin, L., and Wühr, M. (2019). A Review on Quantitative Multiplexed Proteomics. Chembiochem Eur. J. Chem. Biol.

Paulo, J.A., O’Connell, J.D., Everley, R.A., O’Brien, J., Gygi, M.A., and Gygi, S.P. (2016a). Quantitative mass spectrometry-based multiplexing compares the abundance of 5000 S. cerevisiae proteins across 10 carbon sources. J. Proteomics 148, 85–93.

Paulo, J.A., O’Connell, J.D., and Gygi, S.P. (2016b). A Triple Knockout (TKO) Proteomics Standard for Diagnosing Ion Interference in Isobaric Labeling Experiments. J. Am. Soc. Mass Spectrom. 27, 1620–1625.

Peshkin, L., Wühr, M., Pearl, E., Haas, W., Freeman, R.M., Gerhart, J.C., Klein, A.M., Horb, M., Gygi, S.P., and Kirschner, M.W. (2015). On the Relationship of Protein and mRNA Dynamics in Vertebrate Embryonic Development. Dev. Cell 35, 383–394.

Peshkin, L., Gupta, M., Ryazanova, L., and Wuhr, M. (2019). Bayesian Confidence Intervals for Multiplexed Proteomics Integrate Ion-Statistics with Peptide Quantification Concordance. BioRxiv.

Schwanhäusser, B., Busse, D., Li, N., Dittmar, G., Schuchhardt, J., Wolf, J., Chen, W., and Selbach, M. (2011). Global quantification of mammalian gene expression control. Nature 473, 337–342.

Sonnett, M., Gupta, M., Nguyen, T., and Wühr, M. (2018). Quantitative Proteomics for Xenopus Embryos II, Data Analysis. Methods Mol. Biol. Clifton NJ 1865, 195–215.

Wessel, D., and Flügge, U.I. (1984). A method for the quantitative recovery of protein in dilute solution in the presence of detergents and lipids. Anal. Biochem. 138, 141–143.

Wühr, M., Freeman, R.M., Presler, M., Horb, M.E., Peshkin, L., Gygi, S., and Kirschner, M.W. (2014). Deep proteomics of the Xenopus laevis egg using an mRNA-derived reference database. Curr. Biol. CB 24, 1467–1475.

Wühr, M., Güttler, T., Peshkin, L., McAlister, G.C., Sonnett, M., Ishihara, K., Groen, A.C., Presler, M., Erickson, B.K., Mitchison, T.J., et al. (2015). The Nuclear Proteome of a Vertebrate. Curr. Biol. CB.

